# Import and Export of Gold Nanoparticles: Exchange Rate in Cancer Cells and Fibroblasts

**DOI:** 10.1101/092601

**Authors:** Vladimir Ivošev, Gloria Jiménez Sánchez, Darine Abi Haidar, Rana Bazzi, Stéphane Roux, Sandrine Lacombe

## Abstract

Cancer is one of the leading causes of death. Radiation therapy is an important modality used in cancer treatment being highly cost-effective. Major flaw of radiotherapy is lack of selectivity between cancerous and healthy tissues. Amelioration of radiotherapy by using high-Z nanoparticles as radiation enhancers is one of potential solutions. Gold nanoparticles (AuNPs) are commonly used as radioenhancers. Understanding the interaction between cancer cells and AuNPs is essential in order to achieve best possible radioenhancing effects, while sparing healthy tissues. This work aims to elucidate interactions of ultrasmall (core size: 2.4 nm and hydrodynamic diameter (Dh): 4.5 nm) fluorescently labeled AuNPs with various human cell lines. In this perspective we measured uptake dynamics, characterized route of internalization and time of intracellular retention in various cancer cell lines and fibroblasts. Our results show that uptake dynamics and internalization pathways are strongly cell line-dependant. We also demonstrate that higher proportion of internalized nanoparticles resides in cancer cells, compared to fibroblasts, in *in vitro* conditions. This work highlights great complexity of cancerous cells and underlines the necessity for excellent knowledge of biological behaviour for each type of cancer. It also emphasizes the major effort needed for efficient cancer treatments and makes an appeal for further development of highly selective nanoparticles in order to hasten their utilization in clinical conditions.

---

Cancer is main cause of death in most of developed counties, while it is second most common cause of death in majority of developing countires^1^. According to estimates from the International Agency for Research on Cancer (IARC), there were 12.7 million new cancer cases worldwide in 2008 and it is estimated that by 2030 global burden will grow up to 21.4 million new cancer cases and 13.2 million cancer-caused deaths^2^. Currently main cancer therapy treatments are surgery, chemotherapy and radiation therapy. Radiotherapy is an important modality used in treatment of cancer, being highly cost-effective and non-invasive treatment method. Approximately 50% of all cancer patients will receive radiation therapy at certain stage of treatment, with an estimation that radiation therapy contributes approximately 40% towards curative treatment^3,4^.

Radiation therapy with ionizing radiation including X-rays, gamma rays and high-energy particles is employed extensively for treatment of almost all types of solid tumors. Success of radiotherapy depends mainly on the total radiation dose given, with a notion that ionizing radiation does not distinguish cancerous and normal cells. The chemical composition of cancerous and normal cells is too similar for inducing a preferential absorption of radiation exclusively by tumours. Additional drawback of radiotherapy is low radiation tolerance of normal tissues that surround the tumour, limiting extent of irradiation. As a result radiation doses are frequently sub-optimal - in order to limit toxicity, thus impairing efficacy of tumor eradication by radiation therapy. Significant improvements between salubrity and toxicity of treatment have been made with radiotherapy fractionation^5^. The selectivity of radiotherapy can be further improved by using radiosensitizers, designed to strongly absorb ionizing radiation. Since high-Z elements exhibit great propensity to absorb X-ray photons, nanoparticles containing high-Z elements appear as promising radiosensitizers for at least two reasons. First: it is possible to gather a large amount of high-Z elements in nanoparticles in contrast to other molecules; and second: the biodistribution of nanoparticles is better suited for preferential accumulation in solid tumors than in the case of other molecules. Application of tumor-specific nanoparticles in radiotherapy aims to improve the radiation therapy outcomes by inducing more toxicity in tumors and less in normal tissues^6^. Therefore improvement of radiotherapy using nanoparticles rich in high-Z elements as radiation enhancers is a subject of undergoing intense studies^7,8^. Several research groups demonstrated that gold nanoparticles can advantageously combine radiosensitization and X-ray imaging, owing to the presence of gold element in the nanoparticles^9,10^. Among gold nanoparticles, the ultrasmall gadolinium chelate-coated gold nanoparticles (Au@DTDTPA-Gd) received much attention since their biodistribution and preferential accumulation in solid tumors can be monitored by magnetic resonance imaging (MRI) - more suited for clinical tumor imaging than X-ray imaging^11^. Au@DTDTPA-Gd nanoparticles paved the way to MRI-guided radiosensitization in order to improve significantly the selectivity and therefore the efficacy of the radiotherapy^12^. The ultrasmall size of these nanoparticles is an asset for the in vivo application since it allows renal clearance which is a pre-requisite in the case of non-biodegradable nanoparticles, but also for the enhancement of the dose effect. Butterworth *et al.* demonstrated that ultrasmall gold nanoparticles (Au@DTDTPA) exert a higher radiosensitising effect than the large ones^13^. In order to have a clearer idea on the action mode of Au@DTDTPA nanoparticles as radiosensitizers at cellular scale, the study of their internalization is crucial. Improving the knowledge about the dynamics of nanoparticles’ uptake by cancer cells and ensuing effects on radiosensitization are therefore essential^14^. Currently it is difficult to draw general conclusions on production of nanoparticles for optimal cellular uptake, as the rate and mechanism of uptake depends on various biological traits^15^. Furthermore, elucidating the exocytosis and intracellular metabolism of nanoparticles could lead to a better understanding of nanoparticles’ cellular toxicity - i.e., if the nanoparticles are trapped in vesicles and leave the cells intact, they are unlikely to induce cellular toxicity^16^. Solving abovementioned problems will enable highly efficient radiotherapy and the highest gain in radiosensitization by metallic nanoparticles.

Since fluorescence imaging is perfectly adapted for visualizing the occurrence and dynamics of specific particles and biological processes in cancerous cells^17^, Au@DTDTPA nanoparticles were tagged with cyanine 5 (Au@DTDTPA-Cy5 NPs) in order to better understand the uptake, time of intracellular retention and exocytosis efficiency of these NPs in different human cell lines. For achieving abovementioned goals, our study was divided in four interdependent parts focused on: (1) the stability of the dye grafting onto the gold nanoparticles, monitored by fluorescence lifetime microscopy (FLIM); (2) the uptake dynamics of the nanoparticles followed up by flow cytometry and chemical analysis (ICP-OES); (3) the characterization of the pathway of nanoparticles uptake using inhibitors of endocytosis and (4) the rate of exocytosis.

## Results

### FLIM measurements

Fluorescence is extensively studied phenomena where fluorescent molecules emit light after exposure to electromagnetic radiation. Fluorescence lifetime (τ) is average time that electron spends in excited state before returning to ground state^18^. Measurements of fluorescent lifetimes are generally unaffected by variations in the number of fluorophores probed, excitation intensity, collection efficiency and photo-bleaching. In addition to previously mentioned traits, it is known that placing fluorescent molecules near metallic nanostructure leads to enhancement of emission intensity, while reducing emission fluorescence lifetime – phenomenon known as metal-enhanced fluorescence (MEF). MEF is generally attributed to excitation of plasmonic modes within nanostructure and it allows metallic nanoparticle to act as resonant nano-antenna^19^. Consequently it was expected that fluorophore’s fluorescence lifetime decay measured in the immediate vicinity of gold nanoparticle will exhibit a shorter fluorescent lifetime, in contrast to fluorophore alone.

In order to verify abovementioned phenomenon and confirm that fluorophore (Cy5) stays attached to the golden core, we compared fluorescence lifetimes of cancerous cells (U87-MG and HeLa) incubated with Cy5 and incubated with Au@DTDTPA-Cy5 NPs. We measured lifetimes of the two sample groups after 30 minutes and 4 hours of incubation with either Cy5 or Au@DTDTPA-Cy5 NPs, followed by 48 hours of post-incubation period in fresh cell medium. This way we could examine whether there is potential detachment of fluorophore during cellular internalization or in intracellular environment. Figures 1a and 1b summarize results of fluorescent lifetime measurements, showing that fluorescence lifetime of Cy5 was shorter when this near-infrared dye is grafted to Au@DTDTPA nanoparticles in comparison to free Cy5 (τ ≈ 1 ns vs 1.2 ns). This decrease in lifetime, attributed to MEF phenomenon, confirmed the presence of Cy5 in immediate vicinity of gold core. Observed result was valid both after 30 minutes of incubation with Cy5 and NPs (Figure 1a) and after more than 48h of intracellular localization (Figure 1b), with statistical significance of (p<0.01) at both time points. Taken together these results proved that Cy5 in Au@DTDTPA-Cy5 NPs stayed attached to golden core and that we observed fluorescent signal of NPs with fluorophore and not just potentially detached fluorophore alone. This result endorsed experimental procedure for the production of fluorescently tagged nanoparticles and enabled us to perform other measurements.

**Figure 1.**
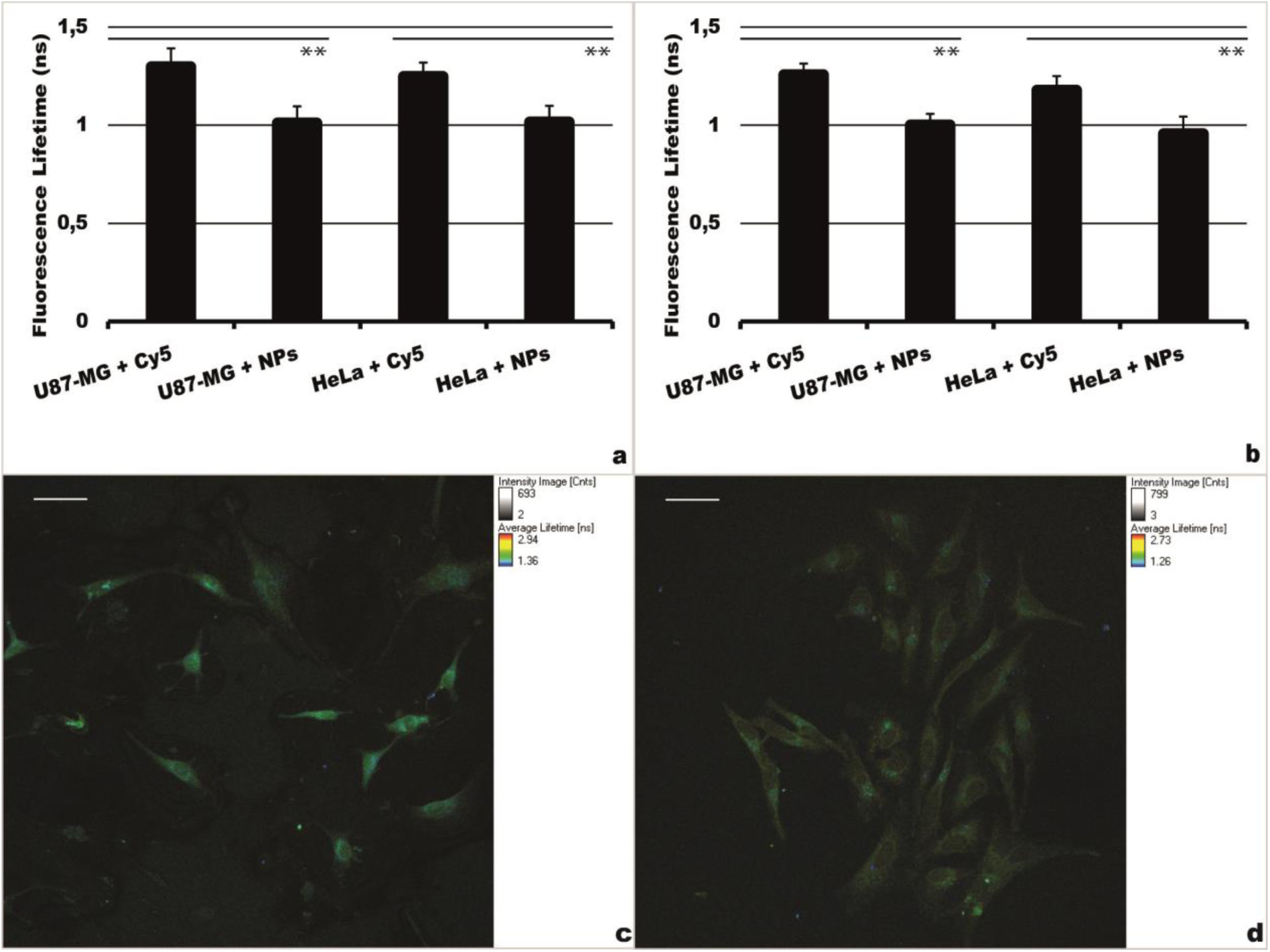
Fluorescence lifetime measurements. Fluorescence lifetime measurements of U87-MG and HeLa cells incubated for **(a)** 30 minutes and **(b)** 4 hours with Cy5 or Au@DTDTPA-Cy5 NPs, followed by 48h of post-incubation in fresh cell media. Images of **(c)** U87-MG cells incubated with Cy5 and **(d)** HeLa cells incubated with Au@DTDTPA-Cy5 NPs. Scale bar is 50 μm. ** under a bar denotes statistically significant difference (p<0.01).

### Uptake dynamics of NPs

FLIM experiments (Figures 1a and 1d) and fluorescence imaging (Figures 1c and 1d) show that fluorescent nanoparticles are internalized by cells without release of organic dye. In other words, the labelling of Au@DTDTPA nanoparticles with Cy5 fluorophores constitutes an efficient strategy for studying the uptake dynamics of these radiosensitising nanoparticles in various human cell lines. Our results showed that there was significant difference among various cell lines both in dynamics of uptake and total amount of uptaken NPs. As Figure 2a shows, even after 30 minutes of incubation with NPs – levels of fluorescence intensity are noticeably higher comparing to cells alone.

**Figure 2.**
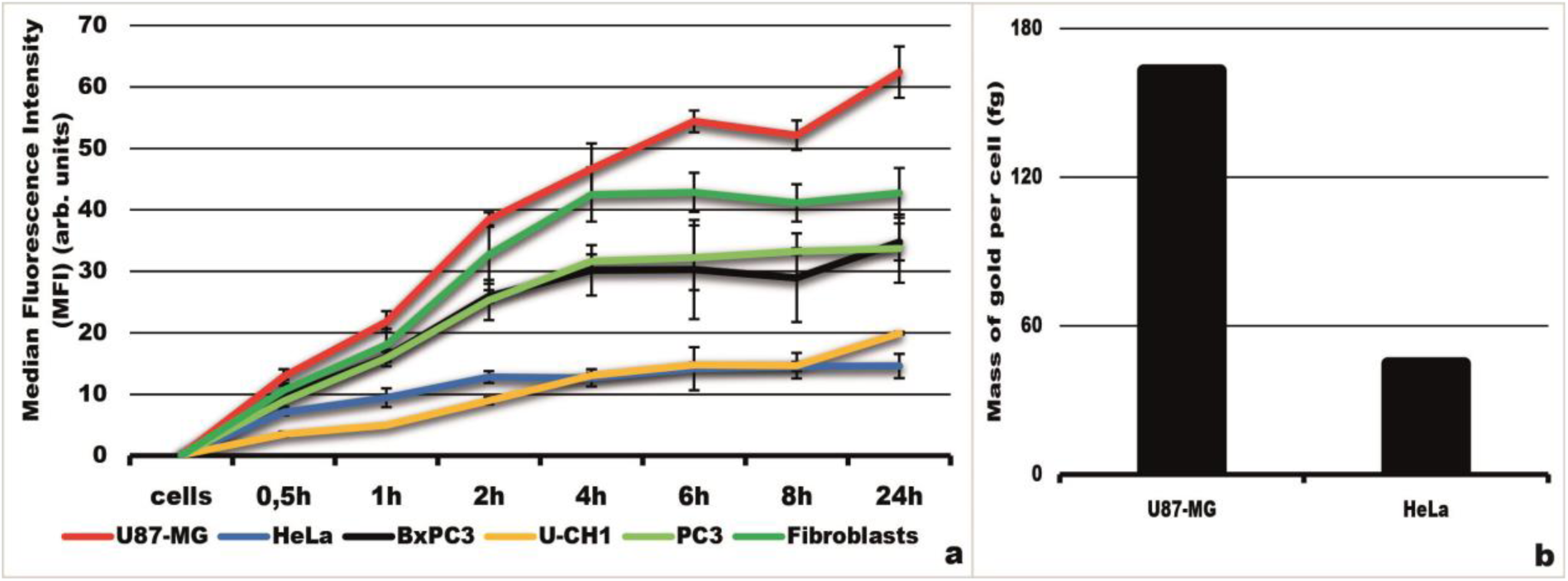
Uptake dynamics of NPs in cancer cells and fibroblasts. **(a)** Uptake dynamics of NPs measured by flow cytometry. Results show mean values of median fluorescence intensity (MFI) of 10 000 cells analyzed in each sample (n=3). Error bars represent standard deviation. **(b)** ICP-OES measurements of intracellular gold content in U87-MG and HeLa cells after 6h of incubation with NPs.

In terms of uptake dynamics we observed three groups of cell lines. U87-MG and human dermal fibroblasts constituted group that speedily engulfed fluorescent Au@DTDTPA-Cy5 NPs. U87-MG and fibroblasts obtained highest uptake levels of NPs as early as 30 minutes of incubation and tended to engulf the highest amounts of NPs at all examined time points. Only difference was that fibroblasts tended to reach plateau of uptake after 4 hours of NPs incubation, while U87-MG cells did not reach their uptake plateau at any of examined time points. Next group consisted of PC-3 and BxPC-3 cell lines that had slightly lower speed of NPs uptake, in comparison with U87-MG and fibroblasts. Just like their rapidly engulfing counterparts (U87-MG and fibroblasts), PC-3 and BxPC-3 cell lines attained their speed of NPs engulfment as early as 30 minutes after incubation with NPs and reached their uptake plateau after 4 hours of incubation (like fibroblasts). Lastly we observed third group of cells, formed by HeLa and U-CH1 cell lines that slowly and inefficiently engulfed NPs - in comparison with previous two groups. HeLa and U-CH1 cell lines also reached their uptake plateau after 4 hours of incubation with NPs. Figure 2a shows that difference in total uptake of NPs was substantial. Comparing cell line with the highest uptake of NPs (U87-MG) and the cell line with the lowest uptake of NPs (HeLa), we detected approximately threefold difference in intracellular content of NPs.

To confirm results obtained by flow cytometry, we performed ICP-OES measurements. We tested cell lines at the end of uptake dynamics spectrum (U87-MG and HeLa cells). We selected 6 hours of cellular incubation with NPs in order to be far from detection threshold limit of the technique as well as to measure levels of intracellular gold after the plateau of internalization has been reached in most of examined cell lines. Furthermore, this time point was of interest due to previous publications of our group, where 6h of cellular incubation with various nanoparticles was considered as optimal^7,20^. Results in Figure 2b show good congruence with results obtained by flow cytometry (Figure 2a). We observed that, on average, HeLa cells uptaked 44.7 femtograms (fg) of gold per cell, while U87-MG cells uptaked 163 fg of gold per cell. ICP-OES measurements validated results obtained by flow cytometry and showed that expected intracellular levels of gold in the rest of examined cell lines were in range between 44.7 fg and 163 fg of gold per cell.

### Characterization of NPs internalization

Internalization of nanoparticles is typically attributed to endocytosis^21^. Endocytic pathways are generally subdivided into: phagocytosis, macropinocytosis, pinocytosis, clathrin-mediated endocytosis, caveolae-mediated endocytosis and lipid raft-mediated endocytosis^22^. Phagocytosis is actin-dependent endocytic process by which cells engulf particles with sizes larger than 500 nm and therefore not of particular interest for this publication. Macropinocytosis is endocytic process by which cells internalize fluids and larger particles together, while pinocytosis is endocytic process by which cells absorb extracellular fluids and small molecules^23^. In addition to abovementioned endocytic pathways that utilize large areas of cellular membrane, clathrin and caveolae-mediated endocytosis are more specific, receptor mediated endocytic pathways, where endocytic vesicles are coated with clathrin and caveolin respectively^24^. Lipid raft-mediated endocytosis occurs by fusion of cholesterol and glycosphingolipid membrane-rich domains and subsequent invagination into intracellular space. Since lipid rafts may contain caveolin, this pathway is closely related to caveolae-mediated endocytosis^25^. In order to precisely distinct the uptake pathways, we performed experiments by using appropriate chemical inhibitors of endocytosis. Detailed description of experiments and concentrations of inhibitors are given in Methods section, all based on bibliographic resources^25,26,27,28^. Main effect of each endocytic inhibitor is reported in Table 1.

**Table 1.** Inhibitors of endocytosis.

| Inhibitor | Effect |
| --- | --- |
| 4°C | Deceleration of metabolism |
| 4°C + Sodium azide (NaN<sub>3</sub>) | Deceleration of metabolism and energy depletion |
| Sodium azide (NaN<sub>3</sub>) | Depletion of intracellular energy |
| Phenylarsine oxide | Inhibition of endocytosis |
| EIPA | Inhibition of macropinocytosis |
| Wortmannin | Inhibition of macropinocytosis |
| Nocodazole | Inhibition of macropinocytosis |
| Chlorpromazine | Inhibition of clathrin-mediated endocytosis |
| Dynasore | Inhibition of clathrin-mediated endocytosis |
| Filipin | Inhibition of caveolae-mediated endocytosis |
| Nystatin | Inhibition of lipid raft formation |
| Cytochalasin D | Disruption of microtubules |
| Monensin | Inhibition of lysosome formation |

Previous work in the field of nanoparticles’ uptake showed that incubation with cells at 4°C prevents intracellular uptake of macromolecules by active pathways and simultaneously quantifies the contribution of simple diffusion^29^. To test this observation we compared levels of NPs uptake at 37°C and 4°C. Therefore, we preincubated all cell lines with medium and NPs at 4°C and compared the results with NPs uptake at 37°C. As Figure 3 shows, uptake of NPs was sensitive to temperature of medium in all cell lines and it was significantly lower at 4°C comparing to 37°C. When endocytosis was verified as main pathway of NPs internalization, we wanted to reveal subtypes of endocytosis that were responsible for NPs internalization.

**Figure 3.**
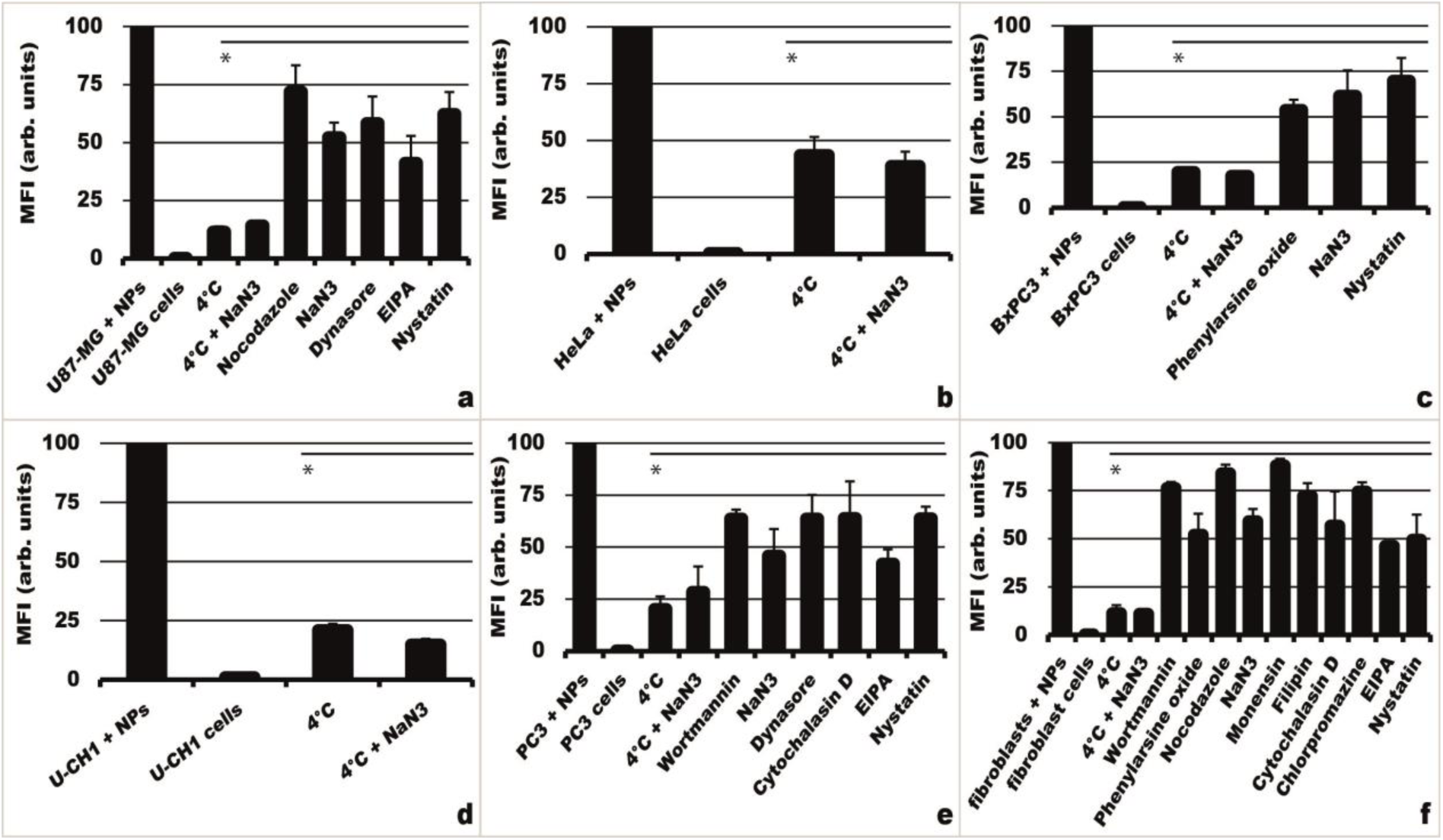
Inhibition of endocytosis in cancer cells and fibroblasts. Results represent normalized values of measured MFI compared with the sample that was not pretreated with any inhibitor of endocytosis. Inhibition of endocytosis in **(a)** U87-MG cells **(b)** HeLa cells **(c)** BxPC3 cells **(d)** U-CH1 cells **(e)** PC3 cells and **(f)** fibroblasts. Results show mean values of median fluorescence intensity of 10 000 cells counted for each sample (n=3). Error bars represent standard deviation. * under a bar denotes statistically significant difference (p<0.05).

As expected from the results in Figure 2a, the predominant pathways of NPs internalization were different among examined cell lines. However, it was possible to observe correlation between the predominant pathway of NPs uptake and observed NPs content in most of cell lines. Figure 3 shows statistically significant effects (p<0.05) of endocytosis inhibitors, while effects of all inhibitors can be found in Supplementary Information (S1). Cells with rapid uptake of NPs (U87-MG and fibroblasts) utilized macropinocytosis as predominant endocytic pathway. This conclusion was substantiated by significant inhibitory effects of EIPA and nocodazole in U87-MG cells, while in case of fibroblasts powerful effects of EIPA, wortmannin and nocodazole were observed. Additionally, lipid rafts played partial role in endocytosis of NPs in fibroblasts, due to inhibitory effect of nystatin on endocytosis. BxPC-3 and PC-3 cell lines engulfed NPs largely through clathrin-mediated endocytosis, although partial contributions of macropinocytosis in PC-3 cell line and lipid rafts in BxPC-3 cell line were observed. Above mentioned conclusions were drawn from substantial effects of phenylarsine oxide and nystatin on endocytosis in BxPC3 cells, while dynasore and cytochalasin D had important role in inhibition of endocytosis in PC-3 cells. Moreover in PC3 cell line micropinocytosis played notable role in engulfment of NPs, due to considerable contributions of EIPA and nystatin on endocytosis inhibition. Lastly, cells with slow and inefficient uptake of NPs, HeLa and U-CH1, were essentially unresponsive to inhibitors of endocytosis. As it is shown in Figures 3b and 3d, majority of endocytosis inhibitors failed to provoke statistically significant inhibition of endocytosis. HeLa cell line showed peculiar behaviour (Figure 3b), where deceleration of metabolism at 4°C did not have enormous effect on endocytic uptake, unlike in the rest of examined cell lines. This indicated that in HeLa cells passive diffusion was an important manner of NPs internalization, even at 37°C.

### Excretion dynamics of NPs

Exocytosis is the process of delivery and fusion of intracellular vesicles with cellular membrane and subsequent extracellular excretion of vesicles content. Exocytosis is diametrically opposite process to endocytosis and it is essential for housekeeping functions of cells^30^. Dynamics of nanoparticles’ exocytosis holds prospective importance for radiotherapy since time of NPs intratumoral retention could play a key role in maximizing radioenhancing effects and optimal treatment planning. Consequently after measuring uptake dynamics and uncovering predominant pathways of NPs uptake, we investigated aptitude of human cell lines to retain engulfed NPs.

Our results showed that dynamics of exocytosis varied among examined cell lines, just like dynamics of endocytosis. We presented exocytosis results as the percentage of maximal measured intracellular NPs content, reached after 4 hours of incubation with NPs at 37°C. Therefore, these results show efficiency of each cell line in retaining internalized NPs and not the absolute levels of internalized NPs - since the aptitude for internalization of NPs varies among cell lines, as shown in Figure 2a. Accordingly Figure 4a shows percentage of intracellular retention of internalized NPs at several time points. As in the case of uptake dynamics measurements, we observed three distinct groups - however cell lines that formed these three groups were different. First group of cell lines (U-CH1 and BxPC-3) retained majority of engulfed NPs, even 48 hours after incubation with NPs was terminated. Interestingly, both U-CH1 and BxPC-3 obtained a plateau of exocytosis 3 hours after incubation with NPs was terminated. Next, we observed a group of cell lines that retained moderate levels of intracellular NPs (HeLa, PC-3 and U87-MG), in comparison with U-CH1 and BxPC-3 cell lines. In case of PC-3 and U87-MG cell lines we noticed higher levels of exocytosis, in comparison with HeLa cell line. Unlike the group that efficiently retained engulfed NPs - PC3, U87-MG and HeLa cell lines retained less than 50% of internalized nanoparticles 48 hours after incubation with NPs was terminated. Lastly we observed third group, where dermal fibroblasts failed to retain engulfed NPs and excreted vast majority of intracellular NPs. Unlike in other examined cell lines, in case of dermal fibroblasts we observed continuous sharp decrease in intracellular concentration of NPs with time, due to high levels of exocytosis in this cell line.

**Figure 4.**
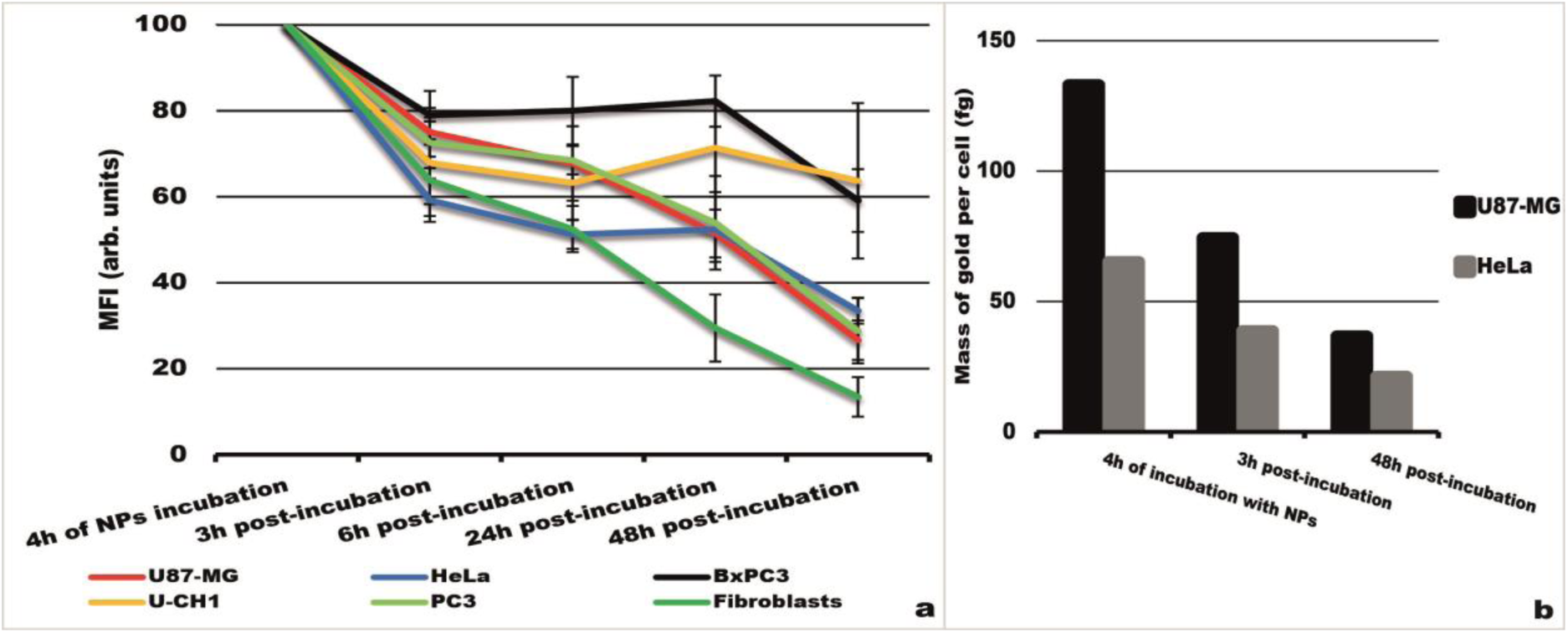
Exocytosis of NPs from cancer cells and fibroblasts. **(a)** Dynamics of exocytosis in cancer cell lines and fibroblasts. Results represent normalized values of measured MFI compared with the sample where cells were incubated with NPs for 4h. Results show mean values of median fluorescence intensity of 10 000 cells counted for each sample (n=3). Error bars represent standard deviation. **(b)** ICP-MS measurements of intracellular gold content. Results show intracellular gold content (fg) in U87-MG and HeLa cells at three time points: after 4h of incubation with NPs; after 4h of incubation with NPs, followed by 3h of post-incubation in fresh cell media; after 4h of incubation with NPs, followed by 48h of post-incubation in fresh cell media.

Observed brisk drop of NPs intracellular content at 48 hours post-incubation time point in all cell lines, except U-CH1, is presumably due to doubling time of U-CH1 cell line (approximately 72 hours) - substantially longer in comparison with the rest of examined cell lines (approximately 24 hours). However that does not invalidate observation that fibroblasts excrete vast majority of internalized NPs, contrary to examined cancer cell lines.

In order to confirm observed trends of NPs exocytosis, we performed ICP-MS experiments. Since we wanted to measure minuscule changes in intracellular gold content, we selected this technique due to its lower detection threshold and higher sensitivity, in comparison with previously used ICP-OES^31^. Thus we compared intracellular levels of NPs in U87-MG and HeLa cell lines at three time points of exocytosis dynamics experiments. As the Figure 4b shows, the levels of intracellular gold content were in accordance with results obtained by flow cytometry. ICP-MS results confirmed sharp drop in concentration of internalized NPs in U87-MG cell line with time and more gradual drop in NPs content in HeLa cell line. ICP-MS results also confirmed that cells excreted substantial amounts of NPs as soon as 3h after the incubation ended. Nevertheless results showed that certain proportion of ingested NPs resided in cells as long as 48 hours after the incubation with cells ended. As expected, the absolute amounts of intracellularly retained NPs were strongly cell line dependant.

## Discussion

This study aimed to measure uptake dynamics, characterize pathways of internalization and measure exocytosis dynamics of 4.5 nm fluorescently tagged Au@DTDTPA-Cy5 nanoparticles in various cancer cell lines and dermal fibroblasts. Results showed that uptake of NPs in most cases reached a plateau after 4 hours of incubation and that total amount of uptaken NPs varied up to threefold - in terms of intracellular gold content among examined cell lines. Afterwards we characterized specific pathways of NPs engulfment and showed that correlation between uptake dynamics and overall aptitude for internalization of NPs can be drawn. Lastly we determined proportion of retained intracellular NPs with time by measuring the exocytosis rates. We observed that levels of intracellular NPs stayed reasonably high for up to 48 hours after incubation, in case of examined cancer cell lines. However we observed that dermal fibroblasts (cells of healthy tissue) excreted internalized NPs rapidly and efficiently, proposing these NPs as a good radioenhancing candidate for radiotherapy.

This work highlighted important considerations for amelioration of radiotherapy with high-Z elements. First of all, it is apparent that aside from differences in uptake of NPs among cell lines, there is substantial heterogeneity in intercellular uptake of NPs in one cell line. This phenomenon is exemplified by relatively large values of standard deviation in most of our results. However, this phenomenon is described in literature and usually attributed to widespread genetic divergence of tumour cells, therefore different biological behaviour, even in the same tumor^32,33^. As the amounts of internalized NPs have direct and proportional significance for radioenhancing effects, the heterogeneity in NPs uptake may play an important role in treatment planning and outcomes of radiation therapy^34,35^.

Important parameter for radiotherapy improvement is the maximal amount of high-Z element that can be engulfed by cancer cells. Due to their size, small NPs (like ones used) have unique advantages over larger nanoparticles in terms of tumour uptake owing to enhanced permeability and retention (EPR) effect^36^. Additionally ultrasmall nanoparticles are preferred for *in vivo* application due to their safer behavior: (1) free circulation in bloodstream, i.e. no uptake by macrophage rich organs (liver, spleen), but accumulation in solid tumors due to EPR effect and (2) body removal by renal clearance - a requirement for non-biodegradable NPs.

Measured by ICP-OES, mass of intracellular gold was in range of 44.7 to 163 fg per individual cell, in HeLa and U87-MG cell lines respectively. Even though intracellular mass of gold is minuscule in comparison with average mass of cancer cell (in order of 1 ng to 3.5 ng^37^), observed intracellular levels can lead to notable radioenhancing effects. Recently published study by Duffort *et al* showed marked radioenhancing effects of nanoparticles even in the concentrations of ppb in cancer tissue^38^. Consequently it is possible to achieve noteworthy radioenhancing effect due to augmentation of deposited nanodoses in cancer cells. It is widely accepted that majority of radioenhancing effects are driven by photoelectric and Compton effects, Auger electrons, photoelectrons and other secondary electrons, reactive oxidative species (ROS) in nanometric distances around internalized NPs^39–41^. Subsequently radioenhancing effect is a result of damage of various cellular organelles like mitochondria, endoplasmic reticulum (ER), Golgi apparatus, etc. and will induce cell death by disrupting cellular homeostasis^42,43^.

Figure 5 summarizes internalization pathways in examined cell lines. Although we used exactly same conditions and NPs in all cell lines, modes of internalization were diverse. This adds another layer of complexity that one has to take into account during production of NPs and radiotherapy treatment planning. It is indispensable to reach most effective compromise between high yield of internalized particles (e.g. by macropinocytosis) and at the same maintaining high targeting specificity for cancerous cells. Equally important is to act at the optimal time, determined by cell-dependant uptake dynamics and dynamics of exocytosis.

**Figure 5.**
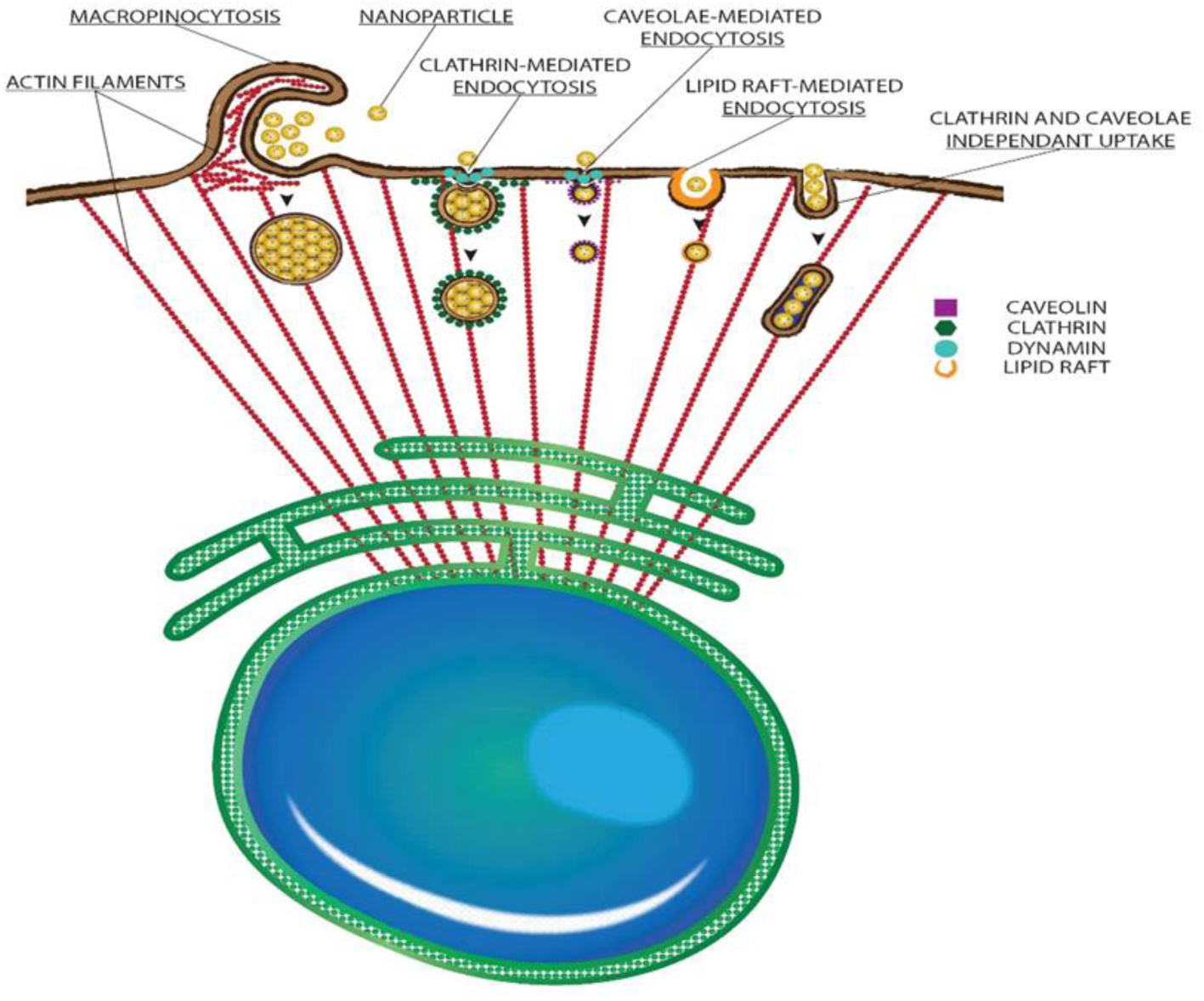
Pathways of nanoparticle internalization. Illustration of observed internalization routes of NPs in examined cancer cells and fibroblasts.

Lastly we showed that levels of internalized NPs in cancer cells remain reasonably high, even 48 hours after incubation with NPs was terminated. On the contrary, we showed that fibroblasts excrete internalized NPs quickly and efficiently. Even though obtained *in vitro*, these results are extremely encouraging for the use of these particular NPs both as imaging agents as well as therapeutic agents. Our observations might be beneficial for radiotherapy, since one can precisely fractionate deposited dose and still achieve radioenhancing effects solely in cancer cells, thus sparing the healthy surrounding tissue. Benefit for potential patients is that it would not be necessary to administer NPs before each radiotherapy session - since the levels of NPs in cancer cells are reasonably high, whereas levels in surrounding healthy tissues are low. As the merit, due to extremely small sizes examined ultrasmall NPs would be safely eliminated through process of renal clearance, without any potential toxicity for liver, spleen and other internal organs of potential patients^44,45^.

In conclusion, this work highlights diversity and complexity of cancer cell responses to a perturbation such as presence of nanoparticles and exemplifies how specific the treatment has to be in order to be maximally effective. Prospective experiments need to determine the exact intracellular localization of NPs in various cell lines and the corresponding effects on radioenhancement and cellular death.

## Methods

### Gold nanoparticles

The synthesis, adapted from the protocol developed by Brust *et al*., consisted of reducing HAuCl_4_·3H_2_O with NaBH4 in presence of thiolated ligands (stabilizers) which, by adsorption on growing particles, ensured control of the core size and the stability of the colloid^46^. In this study, the ligand consisted of a dithiolated derivate of the diethylenetriaminepentaacetic acid (DTPA), named DTDTPA^44^. For the purpose of our experiments AuNPs were tagged with an organic dye, grafted-like Cy5-amine. Stock solution concentration of NPs was 45 mM, while the working concentration was 0.5 mM in all experiments^47^. The concentration of the Cy5 in NPs was 0.007 mM. Zeta potential of NPs was −34 mV at pH 7, measured by Nanosizer ZS (Malvern Technologies). Gold core of NPs has diameter of 2.4 nm, while hydrodynamic diameter in a complete cell medium was 4.48 nm. Hydrodynamic diameter was measured by Viscosizer TD (Malvern Technologies, Saclay, France) and it was an average of four separate measurements.

### Cell lines

All cells were cultivated in standard incubating conditions (37°C, 5% CO_2_) and used for experiments until the 25^th^ generation. For our experiments we used following cell lines: **U87-MG:** human glioblastoma cells extracted from the brain tissue of a 44 year old Caucasian male patient. Their morphology is epithelial and they are adherent in cell culture. **HeLa:** human adenocarcinoma cells extracted from the cervix of 31 year old African female patient. Their morphology is epithelial and they are adherent in cell culture. **PC-3:** human prostate cancer cells derived from the metastatic tissue in the bone of 62 years old Caucasian male patient. Their morphology is epithelial and they are adherent in cell culture. **BxPC-3:** human pancreatic cancer cells derived from the pancreas of the 61 year old female patient. Their morphology is epithelial and they are adherent in cell culture. **U-CH1:** human chordoma cancer cells derived from the sacral bone of the 56 year old Caucasian male patient. Their morphology is mesenchymal-like and they are adherent in cell culture. **Primary Dermal Fibroblasts:** human fibroblasts derived from the foreskin of male African new-born. Their morphology is spindle-shaped and they are adherent in cell culture. All cell lines were bought from ATCC, Manassas, Virginia, USA. U87-MG, HeLa and primary dermal fibroblasts were cultivated in Dulbecco’s Modified Eagle Medium (DMEM) (Life Technologies) supplemented with 10% heat-inactivated fetal bovine serum (PAA), 100 U/ml penicillin (PAA), 100 μg/ml streptomycin (PAA) and 1% Non-Essential Amino Acids (Life Technologies). PC-3 and BxPC3 cells were cultivated in RPMI 1640 (Life Technologies) supplemented with 10% heat-inactivated fetal bovine serum (PAA), 100 U/ml penicillin (PAA) and 100 μg/ml streptomycin (PAA). U-CH1 cells were cultivated in mixture of IMEM (Life Technologies) and RPMI 1640 (Life Technologies) in a ratio of 4:1, supplemented with 10% heat-inactivated fetal bovine serum (PAA), 100 U/ml penicillin (PAA), 100 μg/ml streptomycin (PAA) and 1% Non-Essential Amino Acids (Life Technologies).

### Flow cytometry measurements

Flow-cytometric measurements were performed by Partec CyFlow Space flow cytometer (Partec GmbH, Munster, Germany) with 633 nm excitation from a red solid-state laser. In the flow cytometer, optical filters were set up so that cyanine 5 was measured in the FL7 channel. Data was acquired on two-parameter dot plots of forward scatter (FSC) versus sideward scatter (SSC) by gating cell population of interest. For each sample 10 000 events were selected and analyzed. Data was processed with FlowJo 10 software (FlowJo LLC, Ashland, Oregon, USA).

### Measurements of NPs uptake dynamics

On a day prior to experiment cells were detached from flasks by thoroughly washing with warm (37°C) PBS (2 times) and addition of 0.05% trypsin-EDTA for several minutes. After cells detached from flask, reaction was stopped by adding several milliliters of appropriate media with 10% of fetal bovine serum. Afterwards cell solution was mixed with trypan blue solution in a ratio 1:1 and this mixture was transferred to counting slides of LUNA automated cell counter (Logos Biosystems, South Korea). Cells were counted three times and arithmetic mean was used as final cell number. Then 100 000 cells were transferred to each well of 6-well plate as a single sample. Cells were left overnight and on following day medium was replaced with the fresh one. 0.5 mM of NPs was added to each well for a defined period of time (30 minutes, 1 hour, 2 hours, 4 hours, 6 hours, 8 hours and 24 hours). At the end of incubation period medium with the NPs was removed, each well was washed two times with cold (4°C) PBS and trypsinized with 0.05% trypsin-EDTA for several minutes. Afterwards the reaction was stopped by adding cold complete media (4°C) and the sample was transferred to 15 ml cell culture tubes. Tubes with the cells were centrifuged for 7 minutes at 825 g and the supernatant was decanted from the cell pellet. In next step cells were fixed with 4% paraformaldehyde for 15 minutes. Lastly cold PBS was added to each sample and cells were stored at 4°C until flow cytometry analysis. All the experiments were conducted in minimal presence of light, in order to conserve cyanine 5 from photo bleaching. Samples were analyzed by flow cytometry, as described in previous section.

### Characterization of NPs endocytosis pathways

In order to decipher pathways of endocytosis we used various inhibitors, based on resources in literature (references in Results section). For this purpose we used: 5-(N-Ethyl-N-isopropyl) amiloride (EIPA) [100 μM], chlorpromazine hydrochloride [100 μM], cytochalasin D [40μM], dynasore hydrate [80μM], filipin [1μM], monensin [3μM], nocodazole [50 μM], nystatin [50 μM], phenylarsine oxide [2μM], sodium azide (NaN_3_) [0.1% w/v] and wortmannin [200 nM]. All inhibitors were acquired from Sigma-Aldrich (Merck KGaA, Darmstadt, Germany). Cell were cultivated and prepared for experiments in the same manner as for the uptake dynamics. Experimental procedure only differed in the fact that inhibitors of endocytosis were added to the samples one hour prior to the addition of NPs. After one hour of incubation with inhibitors has passed, the NPs were added for one additional hour. After that samples were prepared in the same manner as for the uptake dynamics experiment and the fluorescence of samples was measured by flow cytometry, as described previously.

### Measurements of NPs intracellular retention times

Cell cultivation and sample preparation was identical as for the studies of uptake dynamics. Only difference was that NPs were incubated with cells for four hours. After four hours of NPs incubation with cells, media with NPs was decanted, cells were washed with warm PBS two times and fresh, complete cell culture medium was added for various time duration (3 hours, 6 hours, 24 hours, and 48 hours). After the time of incubation in fresh medium expired, samples were prepared for the flow cytometry analysis in an identical manner as for the measurements of uptake dynamics.

### Uptake dynamics measurements by ICP–OES

After incubation of adherent cells with NPs for 6 hours, cell samples in complete cell medium were washed two times in PBS, trypsinized for several minutes and centrifuged for seven minutes at 825 g. Subsequently supernatant was decanted and cell pellet conserved in the glass tubes. Prior to the measurements, samples were mineralized with hot ultrapure aqua regia in high purity polypropylene tubes (DigiTUBEs, SCP Science, Courtaboeuf, France). The gold content in the collected cells was determined by ICP-OES using a 710-ES spectrometer (Qualio Laboratories, Besançon, France).

### Excretion dynamics measurements by ICP–MS

After same experimental treatment as for ICP-OES measurements, cell samples in complete cell medium were washed two times in PBS, trypsinized for several minutes and centrifuged for seven minutes at 825 g. Afterwards supernatant was decanted and cell pellet conserved in the glass tubes and stored at −80°C until samples were analysed by ICP-MS (UT2A, Pau, France).

### Fluorescence lifetime measurements

At the IMNC laboratory (Orsay, France), samples were imaged using Leica TCS SP8-FLIM microscope at PIMPA platform. A Mai Tai DeepSee Ti: sapphire laser (Spectra-Physics, Santa Clara, USA) with automated dispersion compensation and a TCS SP8 MP confocal microscope (Leica Microsystems, Wetzlar, Germany) performed fluorescence lifetime imaging of samples under two-photon excitation. The laser cavity had over 2.4 W of average power at 800 nm and was tuneable from 690 nm to 1040 nm. The repetition rate of laser source was 80 MHz and output pulse duration was around 70 fs. Laser was controlled by Leica software, including easy adjustment of the prechirp unit. To collect two-photon fluorescence signal, two super sensitive hybrid detectors (HyD) from Leica were used in non-descanned position. The collected signal passed through a first dichroic transparent to wavelength higher than 680 to cut the laser reflexion, then through a second one (FF495-Di03-25x36) to split the light toward two HyD detectors. Behind each detector an additional filter (FF01-448/20-25, FF01-520/35-25, Semrock, New York, USA) was set to define the detection band. For this study a water-immersion Leica objective was used HCX IRAPO L 25X NA 0.95 and the excitation wavelength was set to 800 nm.

After the acquisition, data was adjusted by a monoexponential fit via Symphotime software (PicoQuant, GmbH, Berlin, Germany) to recover the lifetimes from the measured fluorescence decays.

For each sample, five FLIM images were acquired on different area from the specimen. On each FLIM image, seven region of interest (ROI) were selected on different relevant structures and fitted by a mono exponential decay to extract the fluorescence lifetime value.

The main criteria for an acceptable fit were: (1) a χ2 value less than 1.0 and (2) residuals randomly distributed around 0 within the interval +4 and −4. More detailed explanation about the setup is given by *Zanello et al*^48^. FLIM microscopy was used to visualize the uptake of NPs in cells and to confirm that Cy5 stayed attached to the NPs core throughout entire duration of experiments.

## Acknowledgements

The authors acknowledge financial support from the European Union’s FP7-People Program (Marie Curie Actions) within the Initial Training Network No. 608163 “ARGENT”, the French agency of research (Agence Nationale de la Recherche, ANR 2012 RPIB 0010) and the Région Franche-Comté (PhD grant for GJS). A part of this work was done at PIMPA Platform, partly funded by the French program “Investissement d’Avenir” run by the “Agence Nationale pour la Recherche” (grant “Infrastructure d’avenir en Biologie Santé-ANR-11-INBS-0006”). Authors are thankful to Oliver Nüsse for providing access to flow cytometry facilities and to Sandrine Lécart for providing the access to level 2 laboratory. Lastly the authors thank Sylvaine Linget from Qualio Laboratory for ICP-OES analysis and Melisa Moran-Plaza for help with illustrations.

### Author contributions

V.I. designed and conducted experiments related to uptake dynamics, inhibition of endocytosis and dynamics of exocytosis. He also performed statistical analysis of results and wrote the manuscript together with S.L. and S.R. G. J. S. produced fluorescent gold NPs, under the supervision of R. B. and S. R. G. J. S., R. B. and S. R. analyzed data from ICP-OES measurements. D. A. H. conducted FLIM measurements with V. I. and performed statistical analysis of acquired images. S. L. supervised the experiments and discussed results with V.I. Importantly S. R. and S. L. provided necessary funds for all experiments and measurements. All authors approved content of this publication.

### Additional information

#### Competing financial interests

The authors declare no competing financial interests.

**How to cite this article:** Ivošev, V. et al. Import and export of gold nanoparticles: exchange rate in cancer cells and fibroblasts

